# Characterisation of bacteriophage JD419, a Staphylococcal phage with an unusual morphology and broad host range

**DOI:** 10.1101/2020.11.09.370866

**Authors:** Tingting Feng, Sebastian Leptihn, Ke Dong, Belinda Loh, Yan Zhang, Mingyue Li, Xiaokui Guo, Zelin Cui

**Affiliations:** Department of Laboratory Medicine, Shanghai General Hospital, Shanghai Jiao Tong University School of Medicine, Shanghai 200080, China; Department of Microbiology and Immunology/School of Global Health, Chinese Center for Tropical Diseases Research, Shanghai Jiao Tong University School of Medicine, Shanghai 200025, China; Department of Clinical Pharmacy, Shanghai General Hospital, Shanghai Jiao Tong University School of Medicine, Shanghai 201620, China; Zhejiang University-University of Edinburgh Institute (ZJU-UoE), Zhejiang University, Haining, 314400, China; Department of Pathology and Laboratory Medicine, Perelman School of Medicine, University of Pennsylvania, Philadelphia, PA19104, USA

**Keywords:** *Staphylococcus aureus*, bacteriophage, morphology, genome, host-range

## Abstract

As an antimicrobial therapy, therapeutic phages, also known as “Phage therapy” are able to inactivate multi-drug resistant bacteria such as methicillin and vancomycin resistant *S. aureus* and thus present a possible treatment for infections that are otherwise incurable. In this paper, we present a novel phage called JD419, which has a remarkably wide host-range. The virulent phage JD419 exhibits an elongated capsid and was able to infect and lyse 83 of all 129 tested clinical strains (64.3%) of multi-drug resistant *S. aureus* including MRSA. To evaluate the potential as a therapeutic phage, we tested the ability of phage JD419 to remain infectious after treatment exceeding physiological pH or temperature. The lytic activity of the phage was retained at pH values of 6.0-8.0 and at temperatures below 50°C. As phages sometimes contain virulence genes, we sequenced the complete genome of JD419. The 45509 bp genome contains a predicted 65 ORFs, none of which show homology to any known virulence or antibiotic resistance genes. Our study illustrates that *Staphylococcus* phage JD419 has the potential to be used for diagnostic, prophylaxic and therapeutic purposes.

## Background

While viral diseases such as the SARS-CoV2 pandemic have an immediate effect on human health and activity including travel and trade, antimicrobial resistance (AMR) presents a major threat to mankind which is increasing at a less drastic rate, yet not without dramatic consequences. It is estimated that in 2050 about 10 million people will die annually from untreatable bacterial infections if alternatives and novel antibiotics are not available ^[1]^. The rate at which bacteria gain resistance to available (chemical) antibiotics has been increasing over the decades due to higher availability, but also due to the over- and misuse of antibiotics ^[2]^. While so-called antimicrobial stewardships try to influence the (over-) use of antibiotics in order to decrease the rate of resistance, the SARS-CoV2 pandemic with millions of people infected and hundreds of thousands being treated in hospitals, might exacerbate AMR ^[3]^, as antibiotics are often given prophylactically ^[4]^. Due to this situation, and the lack of industry initiative to develop new antibiotics, alternative strategies have to be developed. One such strategy is the use of bacterial viruses that infect and kill a pathogen, also known as phage therapy ^[5]^ Phage therapy has been used successfully against many Gram-negative and -positive bacteria including *Staphylococcus aureus,* but is not yet a standard therapeutic strategy. *S. aureus* is a particularly concerning pathogen as it exists as a harmless commensal; yet it is able to cause various types of infections, from skin abscesses to lethal blood stream infection, often in hospitals, as one of the nosocomial pathogens. Phage therapy also has great potential for the treatment of methicillin-resistant *S. aureus* as well as of vancomycin-resistant *S. aureus* ^[6–7]^. Several reports have demonstrated that phage therapy could present a promising alternative to combat Staphylococcal infections ^[8–9]^.

While members from other families have been reported, most *Staphylococcus* phages belong to the family of Myoviridae which in some instances exhibit a wide host-range that is considered beneficial for phage therapy ^[10–9]^. *Staphylococcus* phages belonging to the family of Siphoviridae have also been reported worldwide, however many of them display a lysogenic life cycle while sometimes also carrying virulence genes, both undesired characteristics for phage therapy. Several reports have shown the potential use of *Staphylococcus* phages alone or in combination with chemical antibiotics, for the penetration and removal of biofilms ^[11]^ and for the treatment of infections ^[12]^.

The aim of this study was to characterize the so-far undescribed *Staphylococcus* phage JD419 with microbiological and genomic methods to assess its potential for diagnostic, prophylaxis, and therapeutic deployment, in particular for the treatment of *S. aureus* infections. Among other isolated phages, phage JD419 showed a very broad host range which was tested on 129 different clinical *S. aureus* isolates that were multi-drug resistant; almost 65% of all strains could be infected, and thus inactivated, by the phage. We characterized further microbiological parameters, including the adsorption to host strains, measured the infection by monitoring the growth of the host and viral release (lysis), and lastly the stability of the phage to thermal and acid stress. Transmission electron microscopy revealed that phage JD419 is a Siphovirus and displays an elongated capsid characteristic of the family, yet unusually long with a length to width ratio of 2. To ensure that the genome of the phage does not encode any virulence factors including antimicrobial resistance genes, we analysed the complete genome sequence in order to evaluate the safety for deploying JD419 as a therapeutic phage, also describing genes that are related to known phage genes in detail.

## Material and methods

### Bacteria isolates and culture conditions

A total of 129 *S. aureus* isolates were obtained from Ruijin hospital and the sixth people hospital of Shanghai in Shanghai, China. Isolates were grown in liquid LB (Luria-Bertani) medium at 37°C, on solid LB medium (1.5% agar), or in LB soft agar overlays (0.7% agar). The phage JD419 was obtained from a single *S. aureus* isolate during routine culture of isolates in the laboratory of Molecular Microbiology in Shanghai Jiaotong University School of Medicine, Shanghai, China. The strain of *S. aureus* Sa28 isolated from Ruijin Hospital was used for amplification of phage JD419 and used for all subsequent experiments as the standard host strain. In addition, 41 of 129 different strains of *S. aureus* were characterized using MLST and Spa typing method, as previously described ^[13]^.

### *Staphylococcus* phage JD419 amplification and purification

High-titer phage stocks were obtained through amplification in liquid LB medium containing 10 mM MgCl_2_ and 5 uM CaCl_2_. First, *Staphylococcus* phage JD419 was added to host cells of *S. aureus* strain Sa28 at a Multiplicity of Infection (MOI) of 0.01. Then after incubation at 37°C overnight, cell lysis -indicated by increased clarity of the liquid culture-was observed. The lysate was then incubated with chloroform (final concentration was 2%) for 30 min under gentle shaking to kill the residual bacteria and release phage particles from bacterial cells. The entire operation procedure for enrichment and purification of phage particles were conducted as previously ^[14]^. Briefly, the residues of bacteria were removed by centrifugation at 6500 rpm (Beckman, JA18.0), for 15 min. Phages in the supernatant were enriched at 4°C overnight using polyethylene glycol (PEG) 8000 and precipitated (final concentration 10 w/v) by centrifugation at 8500 rpm, for 20 min (Beckman, JA18.0). The pellets were dissolved in TM buffer ((Tris-Mg^2+^ Buffer) 10 mM Tris–HCl (pH 7.5), 100 mM NaCl, 10 mM MgCl_2_, 5 mM CaCl_2_)), vortexed, and PEG8000 was removed after adding the same volume of chloroform and vortexed, centrifuging at 4000 g/min for 10 min. The supernatant contained high concentrations of phage to which CsCl was added (0.5 g/mL). The phages were then purified by discontinuous centrifugation in a CsCl gradient (1.33, 1.45, 1.50, 1.70 g/cm^3^) in TM buffer in Ultra-Clear tubes (Beckman Coulter Inc., Fullerton, CA). Finally, the band containing the enriched phages was removed using a syringe, the sample was then dialyzed against TM buffer and stored at 4°C.

### Electron microscopic imaging

Purified phage particles obtained by dialysis after discontinuous centrifugation in a CsCl gradient (described above), were collected by centrifugation at 33000 × g for 1 h and washed twice in 0.1× PBS (pH 7.4) using a Beckman high-speed centrifuge and a JA-18.1 fixed-angle rotor. The sample was deposited on carbon-coated copper grids and stained with 2% (wt/vol) potassium phosphotungstate (pH7.0). Imaging was performed with a Hitachi H7500 transmission electron microscope (TEM) operating at 80 kV

### Host range of *Staphylococcus* phage JD419

The host range of phage JD419 was analyzed by spotting serial dilutions of the phage on a two-layer soft agar lawn of *S. aureus*. Two microliters of concentrated phage lysate (≈10^8^ PFU/ml) and serial dilutions thereof, were gently pipetted on an Luria-Bertani (LB) medium plate, which was then overlaid with *S. aureus* (OD_600nm_≈0.4) mixed in 0.7% top agar cultured 30 min. The lytic ability of phage isolates was assessed using the clarity of inhibition zone bacterial circles formed. The following scoring system was used: perfectly clear (++++), clear (+++), turbid (++) or faint (+), as published previously ^[14]^. 129 strains of *S. aureus* were used to establish an estimate of the host range of phage JD419.

### Adsorption rate of *Staphylococcus* phage JD419

Phage JD419 was added to *S. aureus* Sa28 at a MOI of 0.0005 and incubated at room temperature without shaking. 100 uL of the mixture of phage JD419 with *S. aureus* Sa28 were collected at 1, 3, 5, 7, 9, 12, 15, 18, 21, 24, 27 and 30 min respectively and centrifuged immediately at 16000g for 30 seconds once collected. Afterwards, the titer of phage JD419 contained in the supernatant was determined by counting the plaques on overlay agar plates. The adsorption rate of phage JD419 to the *S. aureus* Sa28 at the time of i was calculated by the formula of (N_0_-N_i_)/N_0_*100, with N_i_ representing the titer of the phage in the supernatant after co-incubation for i minutes, and N_0_ is the titer of the phage before co-incubation. The proportion of the amount of non-adsorbed phages to the amount of phages used for infection, based on three independent experiments, is shown and standard deviations are indicated.

### One step growth curve of *Staphylococcus* phage JD419

Cells of mid-log *Staphylococcus* phage JD419 were harvested by centrifugation and resuspended in fresh LB medium. Phage JD419 was added to the culture at a MOI of 0.01 and allowed to adsorb for 10 min at room temperature, then incubated at 37 °C with gentle shaking. A sample was taken at the points of 1, 5, 10, 15, 20, 25, 30, 35, 40, 45, 50, 55, 60, 65, 70, 75, 80, 85, 90, 100, and 110 min. After centrifugation, the phage titers were determined by counting plaques of overlay agar plates.

### Burst size of *Staphylococcus* phage JD419

Phage JD419 infected Sa28 at a MOI of 0.01. The burst size of phage JD419 in the host of *Staphylococcus* phage Sa28 was calculated by the formula (Nf-N_0_)/N_0_, where N_f_ represents the titers of phage JD419 at the beginning of next growth circle, and N0 represents the titers of phage JD419 at the beginning of former growth circle.

### Stability of *Staphylococcus* phage JD419

The acid-base stability of phage JD419 was assessed as follows: 100-fold dilutions of the initial phage solution containing 10^8^ PFU/ml, were added to in TM buffer of pH values ranging from 2 to 12 and then incubated at 37°C for 1 hour. Subsequently, the samples were diluted and the titers were determined. To address the thermal stability, phage JD419 was incubated in TM buffer (pH7.5) at different temperatures (24°C, 37°C, 40°C, 50°C, 60°C and 70°C) for 1 h, and then the titers were determined.

### Safety assessment of *Staphylococcus* phage JD419 based on genome sequence

The DNA of *Staphylococcus* phage JD419 was extracted using Aidlab kit (Aidlab Biotechnologies Co.,Ltd, Beijing, China). The complete genome of *Staphylococcus* phage JD419 was sequenced at the Chinese National Human Genome Center in Shanghai. Assembly of quality filtered reads was performed using the platform by 454 Life Sciences Corporation. Nucleotide sequence accession number: GenBank accession MT899504. tRNAs were detected using tRNAscan-SE software ^[15]^. To identify any virulence factors or antibiotic resistant genes in the genome of *Staphylococcus* phage JD419, we performed Blast searches against the antibiotic resistance genes database (ARDB, http://ardb.cbcb.umd.edu/) ^[16]^ and the virulence factors database (http://mvirdb.llnl.gov/; http://www.mgc.ac.cn/VFs/main.htm). The genes with more than 70% coverage and 30% identity were kept as the results. The software PHACTS was used to determine whether *Staphylococcus* phage JD419 was characterized as lytic or lysogenic (template phage) ^[17]^. ORFs were predicted combining the methods of the software glimmer, RAST and GeneMarkS ^[18–20]^.

## Results

### Phage JD419 has a flexible tail attached to a prolate capsid

We analysed the morphology of *Staphylococcus* phage JD419 by employing Transmission Electron Microscopy (TEM). The tail is about 300 nm in length, while the head has a prolate geometry, elongated only in one direction. The length to width ratio is approximately 2, with a width of 50 nm and a length of 100 nm. No tail sheath was observed to be present between the head and tip of the tail. The overall morphology indicates that phage JD419 is a member of the family Siphoviridae in the order Caudovirales, with however an unusually elongated head.

### The adsorption rate of phage JD419 to the host surface is a rapid process

In order to establish how high the binding affinity is, and how reversible the reaction of binding of the phage to its host, we measured the time-dependent adsorption of phages to bacterial cells by simple co-incubation and subsequent counting of phage particles present in the supernatant. After incubation with its host strain Sa28 for 10 min, more than 80% of phages adhered to the bacterial surface, while after 20 min, more than 90% of phages remained attached to the host (Fig.2 A). The observed adsorption rates could be fitted to an exponential decay reaction, with half of the phages attaching to the host cell within approximately 2 1/2 minutes, demonstrating that this reaction is very rapid, in turn indicating that the affinity for the host is likely to be very high.

**Fig.1.**
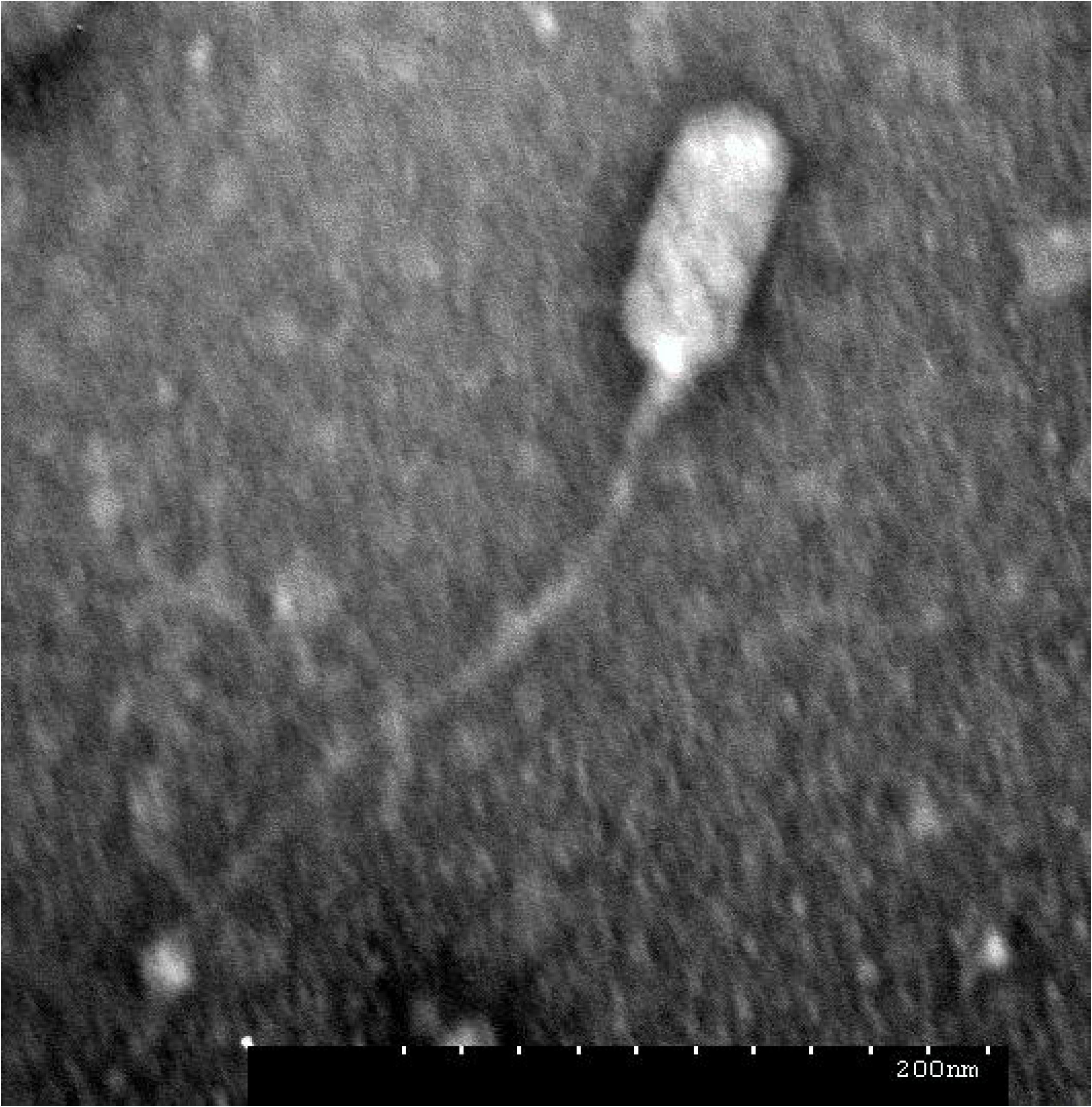
Morphology of *Staphylococcus* phage JD419. The electron micrograph of phage JD419 shows that the tail is flexible and has a length of about 300 nm. The head of the phage is prolate, i.e. extended. The minor axis semi-diameter and major axis semi-diameter of its head are about 50 nm and 100 nm, respecively. Bar: 200 nm.

**Fig.2.**
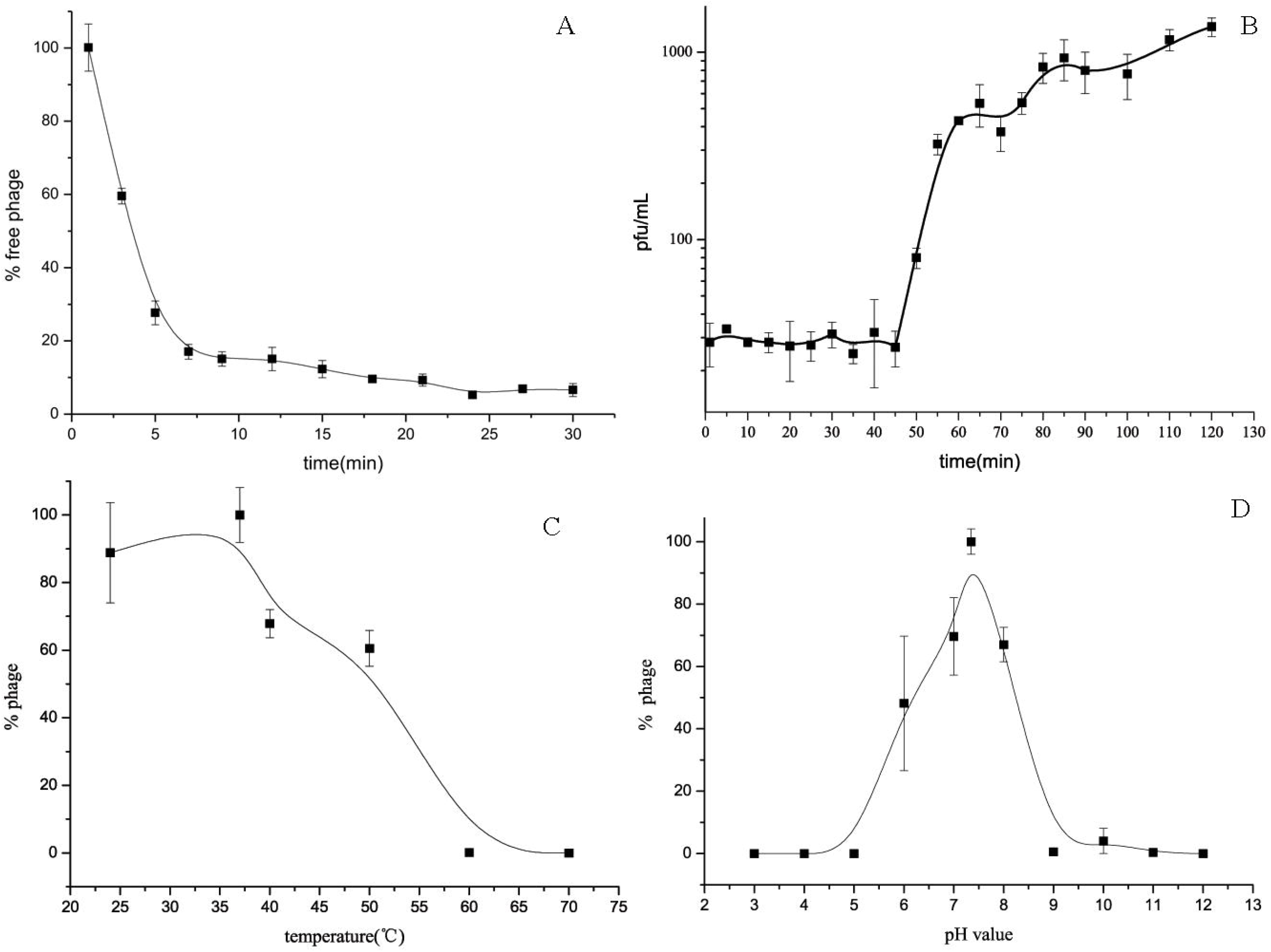
Characterization of *Staphylococcus* phage JD419. (A) Adsorption rate of *Staphylococcus* phage JD419 to its host *S. aureus* Sa28. The X-axis represents the incubation time of *Staphylococcus* phage JD419 and its host, the Y-axis represent the percent of phage that had not been absorbed by its host, error bars represented S.D. (B) One step growth curves of *Staphylococcus* phage JD419; X-axis represents co-culture time of *Staphylococcus* phage JD419 and Sa28, Y-axis represents the change in titer of phage JD419 in the mixture. (C) Thermal stability of phage JD419; X-axis represents different temperatures, Y-axis represents the relative titer of *Staphylococcus* phage JD419 to that the titer at 37°C; % stability = (N/N_0_)×100, where N was the number of viable phages after 1 h of incubation and N_0_ was the initial number of phages. (D) pH stability of *Staphylococcus* phage JD419; X-axis represents the different pH values, Y-axis shows the percentage stability relative to the PFU/mL at pH 7.45.

### Phage replication of JD419 requires more than the typical generation time of the host, releasing about 30 progeny particles

When we infected a growing culture of the *S. aureus* host strain Sa28 with a MOI of 0.01, we could observe the release of phage particles after a latent period of approximately 45 minutes. Complete lysis of all cells was observed at around 70 minutes. The data points of this process were fitted using a sigmoidal fit, showing that half of the phages have been fully released from the cells within 50 minutes. This indicates that the host requires more than one generation time that is commonly observed for its host, *S. aureus*. We also measured the number of phage particles that are being released per cell. Here, we could determine a burst size of approximately 33 phages, those were produced in each cell and subsequently released by lysis.

### Stability of phage JD419 at elevated temperatures and at non-neutral pH values

In order for a phage to potentially be deployed as a therapeutic phage, stability issues have to be considered. We performed experiments to address JD419 stability to acidic or alkaline pH, as well as to elevated temperatures. To test thermal stability, we incubated a phage solution at different temperatures for 1 hour and subsequently determined the phage titer. Compared to 24 °C, the activity, measured as being still an infections particle, only slightly decreased at 40 °C while almost all phages were inactivated when incubated at 60 °C for 1 hour (Fig. 2C). The data points were fitted to a sigmoidal function. The inactivation by change in pH was measured in a range from pH 3 to 12. Phage JD419 was stable at pH values between 6 and 8 (Fig. 2D), making it suitable for most therapeutic application with the exception of the oral route as the phage would be inactivated by the acidic environment of the stomach.

### Phage JD419 exhibits a broad host range

One of the most important parameters for an application as a therapeutic phage is their host range, i.e. how many different bacterial strains the phage is able to infect. We screened 129 different clinical *S. aureus* strains that were obtained from two different hospitals in Shanghai (China). The strains, many of the methicillin -resistant, were classified according to their MLST types, and also included the prevalent types ST239 and ST59. Phage JD419 is capable of lysing 65% of strains tested (83/129). However, the extent of lysis was not identical for all phage-susceptible strains. We categorised the efficiency of the phage according to the plaque appearance which ranged from “completely clear” (++++) to “faint” (+), with “clear” (+++) and “turbid” (++) in between. (Table 2). Most of the phages showed a lysis efficacy of the category “turbid” (++), and “faint” (+), which will require further investigation on the impact of the phage on these particular hosts prior to exploring a potential use of the phage for therapy. Lysis efficiencies were different among isolates of the same MLST type and phage susceptibly did not correlate with MLST types (Table 2).

**Table 1.**
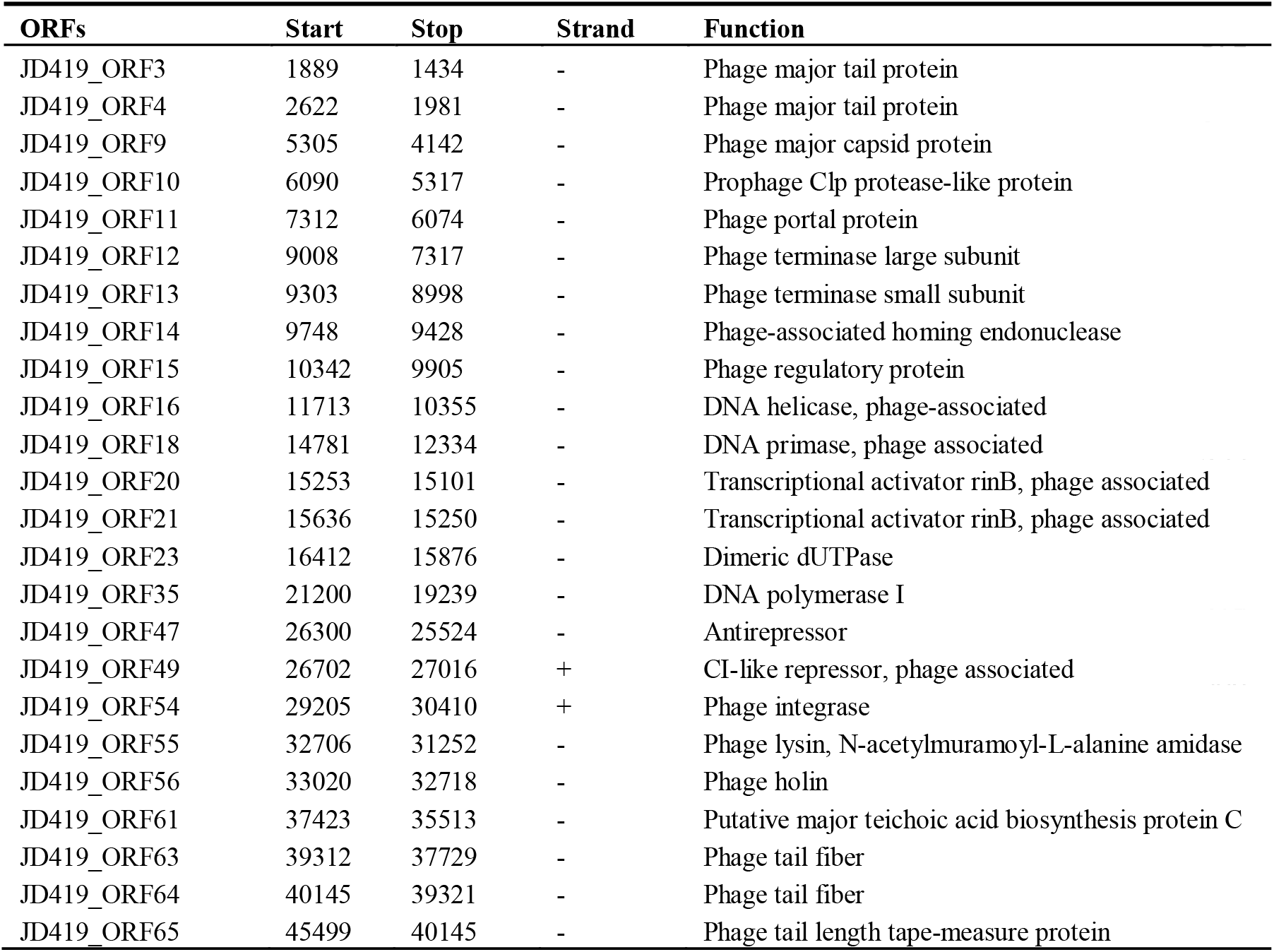
Predicted functional ORFs of *Staphylococcus* phage JD419.

| ORFs | Start | Stop | Strand | Function |
| --- | --- | --- | --- | --- |
| JD419_ORF3 | 1889 | 1434 | - | Phage major tail protein |
| JD419_ORF4 | 2622 | 1981 | - | Phage major tail protein |
| JD419_ORF9 | 5305 | 4142 | - | Phage major capsid protein |
| JD419_ORF10 | 6090 | 5317 | - | Prophage Clp protease-like protein |
| JD419_ORF11 | 7312 | 6074 | - | Phage portal protein |
| JD419_ORF12 | 9008 | 7317 | - | Phage terminase large subunit |
| JD419_ORF13 | 9303 | 8998 | - | Phage terminase small subunit |
| JD419_ORF14 | 9748 | 9428 | - | Phage-associated homing endonuclease |
| JD419_ORF15 | 10342 | 9905 | - | Phage regulatory protein |
| JD419_ORF16 | 11713 | 10355 | - | DNA helicase, phage-associated |
| JD419_ORF18 | 14781 | 12334 | - | DNA primase, phage associated |
| JD419_ORF20 | 15253 | 15101 | - | Transcriptional activator rinB, phage associated |
| JD419_ORF21 | 15636 | 15250 | - | Transcriptional activator rinB, phage associated |
| JD419_ORF23 | 16412 | 15876 | - | Dimeric dUTPase |
| JD419_ORF35 | 21200 | 19239 | - | DNA polymerase I |
| JD419_ORF47 | 26300 | 25524 | - | Antirepressor |
| JD419_ORF49 | 26702 | 27016 | + | CI-like repressor, phage associated |
| JD419_ORF54 | 29205 | 30410 | + | Phage integrase |
| JD419_ORF55 | 32706 | 31252 | - | Phage lysin, N-acetylmuramoyl-L-alanine amidase |
| JD419_ORF56 | 33020 | 32718 | - | Phage holin |
| JD419_ORF61 | 37423 | 35513 | - | Putative major teichoic acid biosynthesis protein C |
| JD419_ORF63 | 39312 | 37729 | - | Phage tail fiber |
| JD419_ORF64 | 40145 | 39321 | - | Phage tail fiber |
| JD419_ORF65 | 45499 | 40145 | - | Phage tail length tape-measure protein |

**Table 2.** Host range of *Staphylococcus* phage JD419.

| *S.aureus* |  |  |  | Phage JD419 |
| --- | --- | --- | --- | --- |
| No. | ST typing | SCCmec | spa |  |
| S130069 | ST1 |  | t127 | +++ |
| S130099 | ST15 |  | t084 | + |
| S130102 | ST1281 |  | t377 | +++ |
| S130174 | ST2315 |  | t11687 | ++ |
| S130060 | ST239 | III | t030 | +++ |
| S130078 | ST239 | III | t037 | +++ |
| S130164 | ST239 | III | t298 | ++++ |
| S130103 | ST239 | III | t459 | +++ |
| S130091 | ST239 |  | t030 | +++ |
| S130084 | ST25 |  | t078 | + |
| S130072 | ST398 |  | t034 | + |
| S130061 | ST398 |  | t1255 | ++++ |
| S130098 | ST398 |  | t571 | + |
| S130076 | ST5 |  | t002 | ++ |
| S130173 | ST5 |  | t570 | + |
| S130065 | ST5 |  | t954 | ++ |
| S130158 | ST59 | IV | t437 | ++ |
| S130064 | ST59 | IV | t441 | + |
| S130177 | ST59 | IV | t163 | +++ |
| S130166 | ST59 |  | t437 | + |
| S130100 | ST6 |  | t701 | ++++ |
| S130105 | ST7 |  | t6248 | ++ |
| S130063 | ST7 |  | t796 | ++ |
| S130070 | ST88 | NT | t12147 | + |
| S130181 | ST965 |  | t062 | +++ |

(Table 2 shows the typing information of some of *S.aureus* that infected by the *Staphylococcus* phage JD419; perfectly clear (++++), clear (+++), turbid (++) or faint (+) )

### Genome analysis of *Staphylococcus* Phage JD419

In order to assess the potential of phage JD419 to be used in phage therapy for treating *S. aureus* infections, we sequenced the complete genome of the bacterial virus. Our analysis showed that the phage has a circular genome with a size of 45509 bp. According to the algorithm of the program PHACTS, JD419 can be characterised as a virulent phage, confirming our laboratory observations. Phage JD419 contains 65 putative open reading frames (ORFs). Most of the proposed ORFs have the same orientation, and several genes overlapped. Putative functions of the ORFs were assigned on the basis of BLASTP analysis and/or domain searches. As shown in Table 1, the ORF encoded proteins can be categorised into five modules which include structure, DNA synthesis, lysis, regulation, and metabolism. Regarding the genes that code for structural proteins, we could identify the major tail protein, the major capsid protein, a portal protein, tail fiber proteins, and a tail length tape-measure protein, among others. ORFs involved in DNA replication included DNA helicase, DNA primase, DNA polymerase I, phage-associated homing endonuclease, phage terminase large subunit, phage terminase small subunit, phage integrase, and finally a phage-associated homing endonuclease. We also were able to identify genes coding for holing and lysin, both crucial for lysis, and a N-acetylmuramoyl-L-alanine amidase. ORFs involved in regulation have also been identified, these including the transcriptional activator rinB, CI-like repressor and a prophage Clp protease-like protein. Lastly, we found ORFs that might possibly influence metabolic processes in the host. These genes coded for example a dimeric dUTPase, and a putative major teichoic acid biosynthesis protein C. While we could confirm that the genome of JD419 does not contain any known virulence factors, we were unable to identify putative roles or functions or a total of 40 of the 65 ORFs encoded proteins, as search algorithms solely reveal that the putative proteins had unknown functions. tRNAs, occasionally found in other large phages such as T4, have not been identified in the genome of JD419.

## Discussion

The clinical use of suitable phages in the treatment of infections might present a potential solution to the threat posed by bacterial pathogens that are antibiotic resistant. Several bacteria are very important, especially in the hospital, as they can cause nosocomial infections, one of which is *Staphylococcus aureus*. The pathogen, which can exist as a harmless commensal bacterium, is able to cause severe and sometimes lethal infections that become increasingly difficult to treat, as many strains have acquired multidrug resistance. Lytic bacteriophages may be deployed aiming to inactivate the pathogen if antibiotics are ineffective as treatment. As in most cases there is no sufficient time to isolate phages that are able to infect the particular strain that causes the infection, it is ideal to have phages that exhibit a broad host range at one’s disposal. In this paper, we characterised the phage JD419 which is able to infect a large percentage of clinical *S. aureus* strains that we tested.

The morphology of JD419 is similar to other reported bacteriophages such as IME-EF1, P70 and 3A ^[21–22]^. However, in general, phages with such massively elongated capsids and a flexible tail have not been described much in the literature. Interestingly, the above mentioned three phages with similar morphology infect only Gram-positive bacteria (Table 3). The phage IME-EF1 infects strains of *Enterococcus feacalís* ^[22]^, phage P70 infects *Lístería monocytogenes* ^[21]^, and phage 3A *S. aureus* (NC_007053.1). While it might be fair to assume that the latter is related to JD419, we were surprised to find their identity to have only 50% similarity. Determining the genome sequence allowed not only a comparison with the phage 3A but also to search for potentially contraindicative factors such as genes coding for virulence or antibiotic resistance. Bacteriophages, in particular lysogenic ones, can be seen as vectors for disseminating antibiotic resistance or virulence genes, as the genome of phages are able to integrate into bacterial chromosome and become prophages.

**Table 3.** Sightings of phages with similar morphology to *Staphylococcus* phage JD419.

| Phage | Host | Head diameter (nm) | Tail length (nm) | References and comments |
| --- | --- | --- | --- | --- |
| IME-EF1 | *Enterococcus faecalis* |  |  | [22] |
| P70 | *Listeria monocytogenes* |  |  | [21] |
| 3A | *S. aureus* | 50×100 | 350 | http://www.phage.ulaval.ca |
| JD419 | *S. aureus* |  |  | This study |

As such phages might bring the potential risk of increasing virulence of the target strains or even introducing antibiotic resistance, the deployment of lysogenic phages in phage therapy should be avoided ^[10]^. Therefore, it is essential to analyze the genome of a phage before using them in clinical applications ^[10]^. In the 45kb genome of JD419, we were able to identify 65 ORFs, none of which encode for any known virulence gene or genes that confer antibiotic resistance. 41 ORFs have unknown functions to date. While the remaining ORFs show similarity to 1) genes involved in the structure of the bacteriophage (major tail protein, major capsid protein, portal protein, tail fiber, and tail length tape-measure protein), 2) genes that are responsible for phage packaging (phage portal protein, phage terminase large subunit and phage terminase small subunit), or 3) genes that allow the release of phage progeny from the host cell (lysin, N-acetylmuramoyl-L-alanine amidase (EC 3.5.1.28), and holin). In addition, we could identify ORFs involved DNA manipulation, including an exonuclease, a DNA helicase, a phage associated DNA primase, DNA polymerase I, a gene with homology to a phage integrase. While the latter gene might indicate that the phage has the potential to integrate into the genome of the host (i. e. being lysogenic) as integrases facilitate the insertion of the viral genome into the bacterial chromosome to become a prophage. Two ORFs encode the transcriptional activator rinB, which might regulate the replication of the phage while another gene (ORF47) encoded an anti-repressor and ORF49 encodes for a CI-like repressor, which shows homology to a prophage repressor, with 99% similarity ^[23]^. Many ORFs are of unknown function but show, in many cases, similarities to ORFs encoded by other phages such as the *Staphylococcal* phage JD007 that however belongs to the *Myoviridae* ^[24]^.

Our microbiological and biochemical characterisation demonstrated the potential of the phage to be used for phage therapy: JD419 has a very wide host-range which was tested on clinical strains of *S. aureus* isolated from patients in Chinese hospitals. Phage JD419 is capable of infecting around 65% of clinical *S. aureus* strains which belong to various MLST types. It has to be stated that the bactericidal activity towards different strains varied significantly, and further experiments are required to establish the “killing potential” of JD419, as from our studies we can only conclude that the phage can infect and perturb the growth of the strains but not if lysis is a fast and efficient process. Therefore, phage JD419’s bactericidal activity towards the target *S. aureus* strain causing the infection would have to be confirmed prior to using it for therapy ^[25]^. The general disadvantage of phages being effective only on a limited number of strains can be overcome by establishing a phage library containing different kinds of phages that cover most strains, of which JD419 could be part of ^[26]^. A highly efficient phage or a so-called phage cocktail of efficient lytic phages is considered optimal for phage therapy. In addition, phages should show some degree of stability to withstand preparatory processes during the development of clinical solutions or in formulations for subsequent patient administration. Phage JD419 exhibits a suitable degree of stability towards pH and temperature, indicating that the phage might be a good candidate as a therapeutic phage. The lability towards acidic pH values indicates that the phage might need to be encapsulated if it was to be delivered through the oral route; however, *S. aureus* infections are usually observed in environments which display a pH close to neutral, easily tolerated by JD419, such as in the epidermis or in the blood.

**Fig.3.**
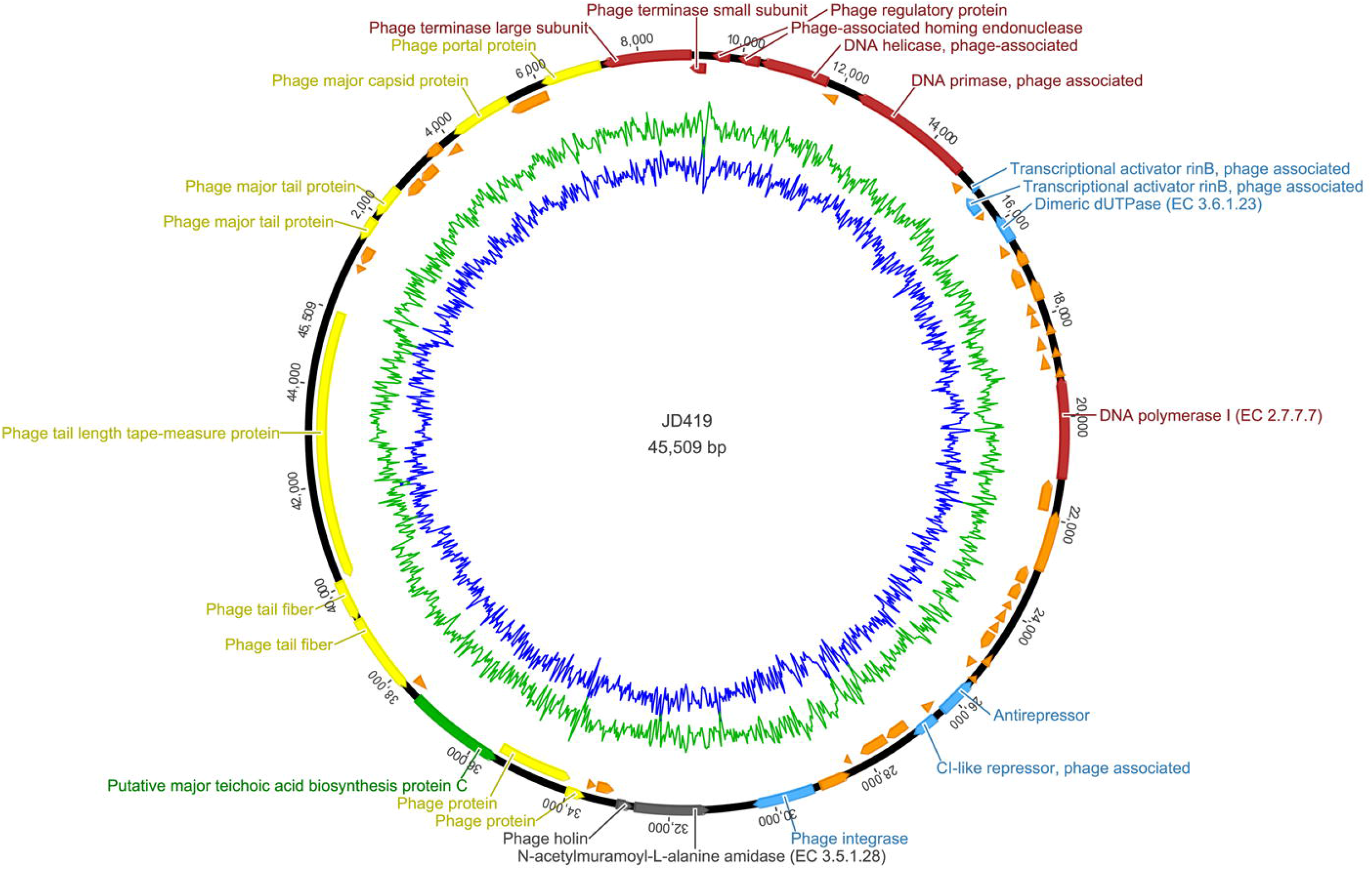
Complete genome sequence of *Staphylococcus* phage JD419. Genome map showing predicted ORFs. Arrows indicated the direction of transcription. Proposed functional modules have been marked based on BLAST and domain search results.

## Author Contributions

Conceived and designed the experiments: Zelin Cui and Xiaokui Guo. Performed the experiments: Zelin Cui and Tingting Feng. Analyzed the data: Tingting Feng, Belinda Loh, Sebastian Leptihn, Mingyue Li and Zelin Cui, Contributed reagents/materials/analysis tools: Zelin Cui, Ke Dong, Belinda Loh, Tingting Feng, and Yan Zhang. Wrote the paper: Tingting Feng, Belinda Loh, Sebastian Leptihn and Zelin Cui.

## Acknowledgements

This work was sponsored by this work was sponsored by the outstanding medical youth program A of Shanghai General Hospital (No.06N1702002), and the National Natural Science Foundation of China (No. 81872540 and 31500154).

